# An Iterative Approach to Polish the Nanopore Sequencing Basecalling for Therapeutic RNA Quality Control

**DOI:** 10.1101/2024.09.12.612711

**Authors:** Ziyuan Wang, Mei-Juan Tu, Ziyang Liu, Katherine K. Wang, Yinshan Fang, Ning Hao, Hao Helen Zhang, Jianwen Que, Xiaoxiao Sun, Ai-Ming Yu, Hongxu Ding

## Abstract

Nucleotide modifications deviate nanopore sequencing readouts, therefore generating artifacts during the basecalling of sequence backbones. Here, we present an iterative approach to polish modification-disturbed basecalling results. We show such an approach is able to promote the basecalling accuracy of both artificially-synthesized and real-world molecules. With demonstrated efficacy and reliability, we exploit the approach to precisely basecall therapeutic RNAs consisting of artificial or natural modifications, as the basis for quantifying the purity and integrity of vaccine mRNAs which are transcribed *in vitro*, and for determining modification hotspots of novel therapeutic RNA interference (RNAi) molecules which are bioengineered (BioRNA) *in vivo*.

## INTRODUCTION

In nanopore sequencing bioinformatics, basecalling refers to the process of interpreting nucleotide sequence backbones from ionic current signals^1^. State-of-the-art basecalling practices in general rely upon deep learning algorithms, e.g. Guppy, Bonito and Dorado provided by Oxford Nanopore Technologies. Previous studies^2-8^ have confirmed that the accuracy of these basecallers is highly likely to be compromised by chemically modified nucleotides, which commonly exist within native DNA and RNA molecules. For instance, precisely basecalling modification-rich tRNAs has long been a challenge. According to a most recent study in yeast, basecalling artifacts were frequently-found near modification hotspots, thereby introducing excessive alignment mismatches, deletions and insertions as opposed to the reference genome^9^. Such a compromised basecalling thus hinder the rigorous investigation of nucleotide sequences using nanopore sequencing.

More recently, nanopore sequencing has been used for quality control^10^ and optimizing^11^ therapeutic RNAs. For example, mRNA vaccines produced through *in vitro* transcription are usually densely-modified (e.g. ubiquitously substituting uridines with pseudouridines or N1-methylpseudouridines), to reduce innate immune responses and promote stability and efficacy^12^. Such dense modifications might strongly compromise basecalling, further biasing the quantification of mRNA vaccine purity and integrity. As reported by the latest quality control study, modification-disturbed basecalling resulted in the overestimation of truncated (therefore low-quality) vaccine mRNAs, compared to the ground-truth capillary electrophoresis result^10^. Consequently, the validity of nanopore sequencing as a generic method to characterize therapeutic RNA was significantly undermined.

To address such bioinformatic challenges, we present an iterative workflow for polishing the nanopore sequencing basecalling. This iterative basecalling method builds upon the hypothesis that initial analyses with, e.g. Guppy, Bonito and Dorado produce acceptable basecalling accuracy. Iterative basecalling next polishes such yielded sketch sequences by aligning them to ground-truth references. Such polished sequences, paired with their raw nanopore sequencing signals, will be used to train the updated basecallers. Such a process iterates until basecalling accuracy converges, as shown in Figure 1A.

**Figure 1.**
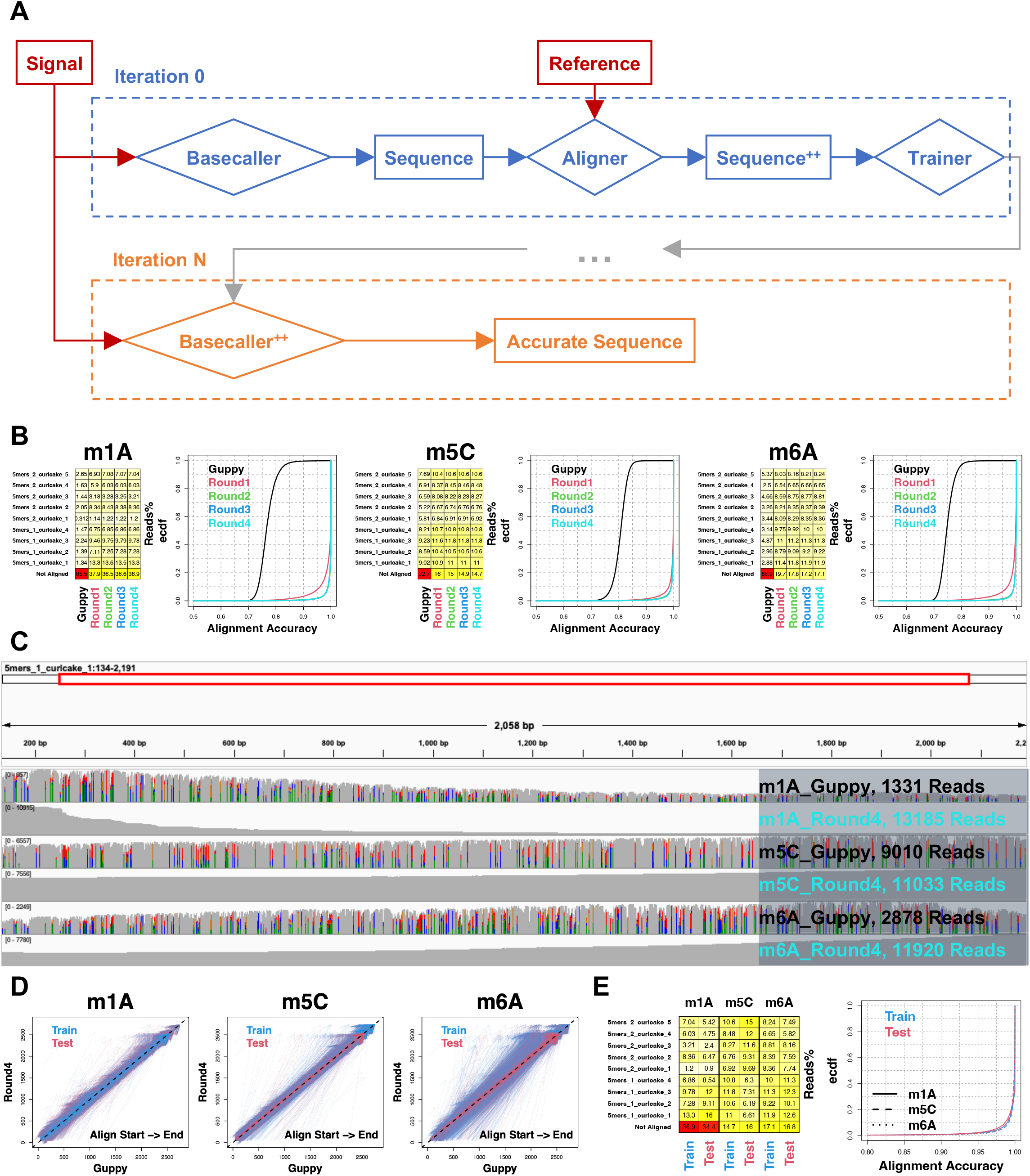
Polishing nanopore sequencing basecalling using an iterative approach. (A) Workflow overview. (B) Mappability and alignment accuracy of RNA control oligos, in which counterparting canonical nucleotides were substituted using N1-methyladenosine (m1A), N6-methyladenosine (m6A) or 5-methylcytosine (m5C) respectively. Distributions of alignment accuracy were shown with ecdf (empirical cumulative distribution function). Guppy, results produced by the Guppy basecaller; Round1-4, results produced by each iteration. (C) RNA control oligo alignments from Guppy and iterative basecalling Round4 were visualized with IGV. Without losing generality, reads aligned to the reference contig “5mers_1_curlcake_1” were shown. (D) Alignment position comparison between Guppy and iterative basecalling Round4 results. Train, results for train datasets used in panel B and C; Test, results for test datasets that are independent of “Train”. Arrows represent alignment directions. (E) Mappability and alignment accuracy comparison between train and test datasets.

## RESULTS

### Iterative basecalling for accurately decoding nucleotide sequence backbones

We benchmarked iterative basecalling with RNA control oligos^8^. Among these oligos, all the counterparting canonical nucleotides were substituted with N1-methyladenosine (m1A), N6-methyladenosine (m6A) or 5-methylcytosine (m5C). We first basecalled and aligned these oligos using Guppy, and found compromised mappability and alignment accuracy. We next executed iterative basecalling, and noticed significant improvements and quick convergence in terms of basecalling performance (Figure 1B, see METHODS). We then confirmed the improvement of mappability and clearance of basecalling artifacts through iterative basecalling with IGV (Figure 1C).

We also confirm that our iterative approach will polish, rather than deviating basecalling. We ran end-product basecallers that were trained in Figure 1B and Guppy on both train and independent test datasets. By comparing yielded sequences, we noticed consistent alignment positions (Figure 1D). Such results confirm that iterative basecalling produces the same sequence scaffolds, with higher per-nucleotide accuracy, compared to Guppy. We further confirmed that iterative basecalling will decode actual nucleotide sequences, rather than overfitting sketch sequences initially basecalled using Guppy. By comparing mappability and alignment accuracy between train and test datasets, we only observed negligible differences (Figure 1E).

### rRNA and tRNA sequence analysis

We applied iterative basecalling to analyze RNAs in real-world biological scenarios. As a proof-of-concept, we first analyzed 18S and 25S yeast rRNAs 1) purified from mutant strains (CBF5_GLU and NOP58_GLU) which have distinct modification patterns; 2) produced using *in vitro* transcription (IVT) which contain no modification^13^. We iteratively trained then benchmarked a basecaller for yeast rRNAs (Figure S1A and B, see METHODS), further applying it on the independent test dataset. Iterative basecalling could still improve mappability, even though great performance has already been made by Guppy. Meanwhile, iterative basecalling can significantly remove basecalling errors, thus increasing alignment accuracy. In addition, we observed similar mappability and alignment accuracy between train and test datasets, which suggested a negligible overfitting when training the iterative basecaller. Noticeably, we observed that the basecalling of IVT rRNAs, which contain no modification, could also be optimized by iterative basecalling (Figure 2A and B, see METHODS). Such an observation suggested that the “vanilla” Guppy basecaller might be suboptimal to analyze certain RNA species, which could be accurately analyzed using iterative basecalling instead.

**Figure 2.**
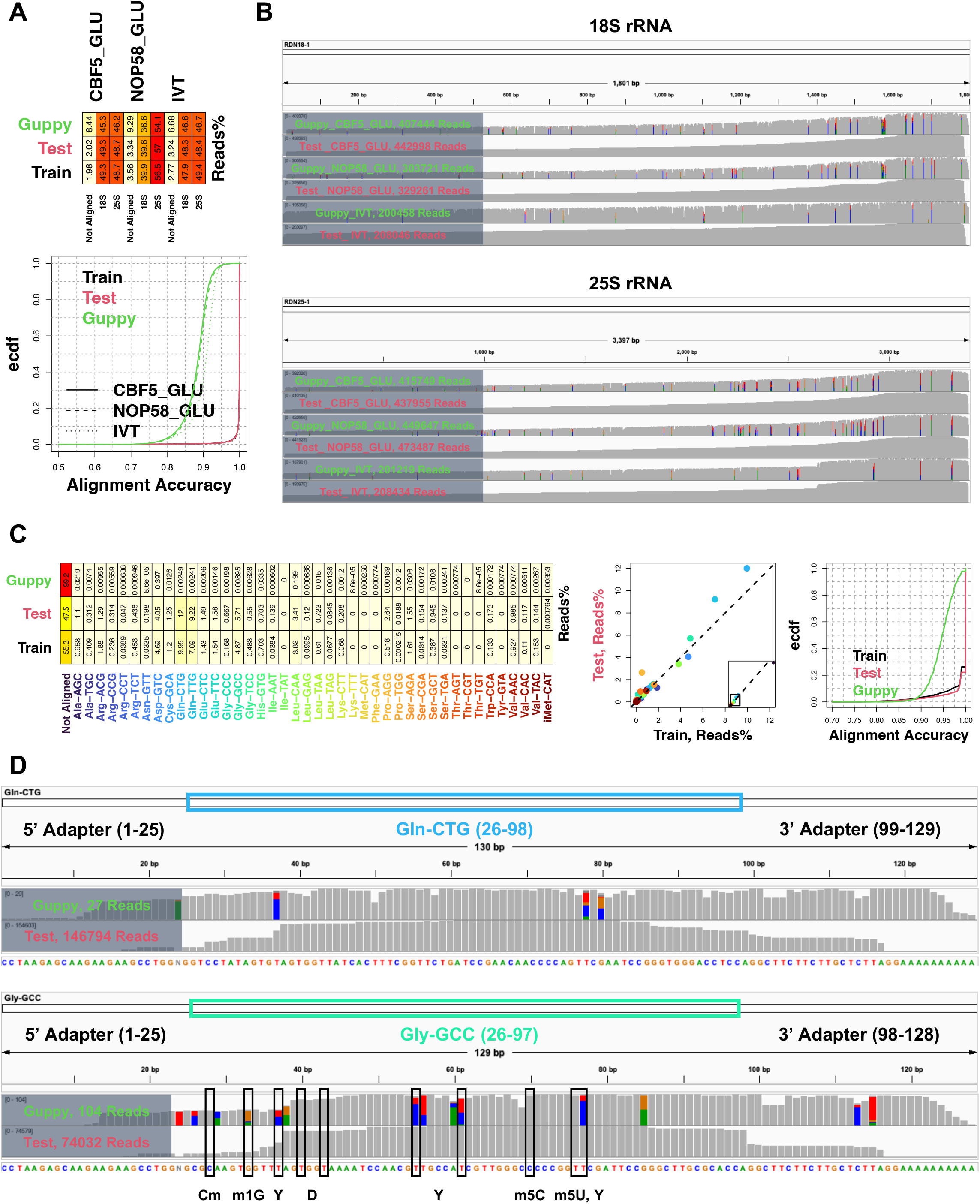
Iterative basecalling improves yeast rRNA and tRNA sequence analysis. (A) Mappability and alignment accuracy of yeast rRNAs. CBF5_GLU and NOP58_GLU, mutant strain rRNAs which have different RNA modification patterns; IVT, rRNAs made by *in vitro* transcription which contain no modification. Train, polished basecalling results for datasets used during iterative basecaller training; Test, polished basecalling results for independent test datasets; Guppy, initial Guppy basecalling results for test datasets. IGV visualization for initial Guppy and polished basecalling results. (C) Mappability and alignment accuracy of native yeast tRNAs. Train, polished basecalling results for datasets used during iterative basecaller training; Test, polished basecalling results for independent test datasets; Guppy, initial Guppy basecalling results for test datasets. (D) IGV visualization for initial Guppy and polished basecalling results. Yeast tRNA species Gln-CTG and Gly-GCC were chosen for visualization, as examples for all-canonical and modification-containing transcripts, respectively.

We also analyzed nano-tRNAseq profiles, which surveyed 42 yeast tRNA species, each having unique modification patterns ranging from densely-modified to all-canonical^9^. We trained then benchmarked the iterative basecaller (Figure S1C and D, see METHODS), further applied it on an independent biological replicate test dataset. As shown in Figure 2C and D, similar to yeast rRNA analyses, we noticed remarkably improved basecalling over Guppy, consistent performance between train and test reads, as well as optimized basecalling of all-canonical molecules such as tRNA Gln-CTG (see METHODS). Taken together, we proved the precise sequence interpretation of biological RNA molecules by the iterative basecalling workflow.

### mRNA vaccine purity and integrity analysis

Building upon the benchmarked efficacy and reliability of iterative basecalling, we next leveraged such a workflow to quantify the purity and integrity of vaccine mRNAs. As a proof-of-concept, we revisited datasets and conclusions from the VAX-seq study^10^. VAX-seq used a plasmid-based system to *in vitro* transcribe vaccine mRNAs, then exploited sequencing and bioinformatic approaches for the corresponding quality control. Without losing generality, VAX-seq was demonstrated using eGFP as the “mock vaccine mRNA”. In particular, VAX-seq performed native RNA nanopore sequencing on all-canonical and U-to-N1-methylpseudouridines fully-replaced mRNA vaccines, and concluded that modifications yielded a significantly higher fraction of truncated transcripts. Contradictorily, VAX-seq reported only a minor transcript length difference between all-canonical and fully-replaced mRNAs by capillary electrophoresis. To scrutinize such a discrepancy, we performed iterative basecalling (Figure S2A and B, see METHODS), and observed significant and consistent basecalling improvements on both train and test datasets (Figure 3A and B, see METHODS). We then determined the alignment status breakdown for test reads, and noticed no reverse, as well as negligible non-primary and supplementary alignments (Figure 3C). We further surveyed alignment positions, and noticed no off-target (overlapping with the plasmid backbone), and >80% full-length (spanning the entire eGFP coding sequence) transcripts, in both all-canonical and fully-replaced samples (Figure 3D). Altogether, such results demonstrated the purity and integrity of N1-methylpseudouridylated mRNA vaccines produced with the VAX-seq plasmid-based *in vitro* transcription system.

**Figure 3.**
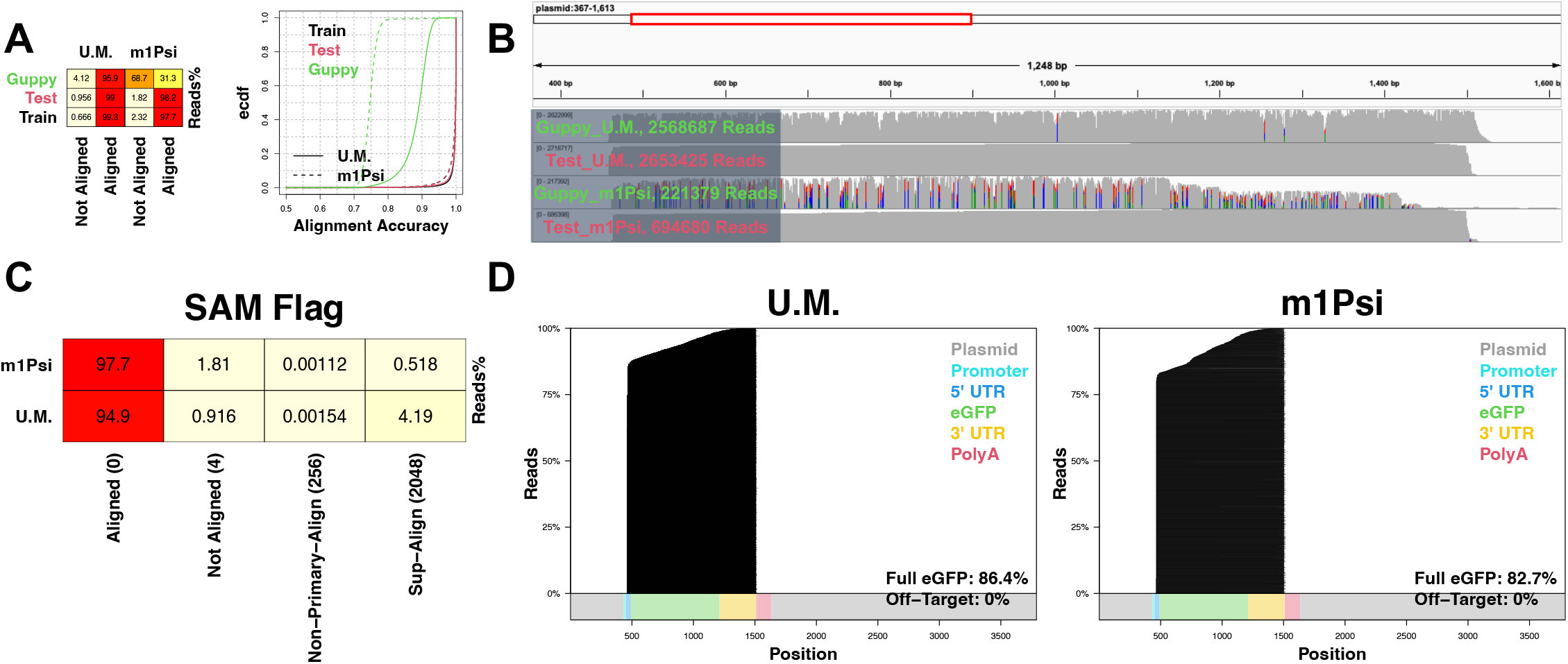
Iterative basecalling determines mRNA vaccine purity and integrity. (A) Mappability and alignment accuracy of vaccine mRNAs. U.M., canonical mRNA; m1Psi, U-to-N1-methylpseudouridines fully-replaced mRNA. Train, polished basecalling results for datasets used during iterative basecaller training; Test, polished basecalling results for independent test datasets; Guppy, initial Guppy basecalling results for test datasets. (B) IGV visualization for initial Guppy and polished basecalling results. (C) The SAM file flag breakdowns, which encode alignment status further representing vaccine purity, for test reads. (D) Per-read alignments, which represent vaccine integrity, for test reads.

### BioRNA modification hotspot inspection

Precisely resolved sequence backbones by the iterative basecalling enhance the rigorous determination of nucleotide modifications. As a real-world preclinical usecase, we inspected modification hotspots within BioRNAs. BioRNAs are engineered from human tRNAs, and are molecular carriers for therapeutic RNA interference (RNAi) agents. In particular, the candidate RNAi agent is incorporated in the BioRNA anti-codon loop for the stable large-scale production, and released by the endogenous microRNA processing pathway after the cellular absorption^14^. Our previous effort suggested the indispensability of modification hotspots for the BioRNA folding and metabolic stability^15,16^. Therefore, we controlled the BioRNA quality from the modification perspective. We prioritized two widely-used BioRNA carriers (BioRNA^Ser^ and BioRNA^Leu^) in our analysis. In these carriers, we used the Sephadex aptamer as the placeholder for therapeutic RNAi molecules. In addition, we synthesized and sequenced their canonical counterparts (ChemoRNA^Ser^ and ChemoRNA^Leu^) as control samples (see METHODS).

For each Bio/ChemoRNA category, we iteratively trained and benchmarked a basecaller (Figure S3A and B, see METHODS), further applying it on the independent test dataset. We found that iterative basecalling could generate significant and consistent basecalling improvements, even in unmodified ChemoRNAs, on both train and test datasets (Figure 4A and B, see METHODS). Based on these precisely determined sequence backbones, we inspected BioRNA modification hotspots. Specifically, we calculated the signal mean distribution for every nucleotide using Remora, then quantified the distribution difference between BioRNA and ChemoRNA using the KS-test D-value (see METHODS). A higher D-value represents a stronger BioRNA signal deviation, which suggests the presence of modification. As shown in Figure 4C, we discovered potential modification hotspots near Positions 19, 71 and 98, for both BioRNA^Ser^ and BioRNA^Leu^. These modification hotspots are located in D, anti-codon and T-arms respectively, and are conserved across different human tRNA species. The preservation of these hotspots therefore indicated the correct folding and metabolic stability of our BioRNAs. Meanwhile, we detected another hotspot (Position 8) in the acceptor-stem. Such a discovery contradicts our prior knowledge that the Position 8 is unmodified in human tRNAs. Considering BioRNAs are produced using *E*.*coli*, and the *E*.*coli* tRNA U8 (U at Position 8) is in general converted into 4-thiouridine (s4U) through enzymatic reactions^17^, we explained this hotspot as an *E*.*coli*-specific s4U modification. For this hotspot, further characterizations on its biochemical and molecular properties, as well as investigations into its influence on the BioRNA therapeutic efficacy are planned as our future efforts.

**Figure 4.**
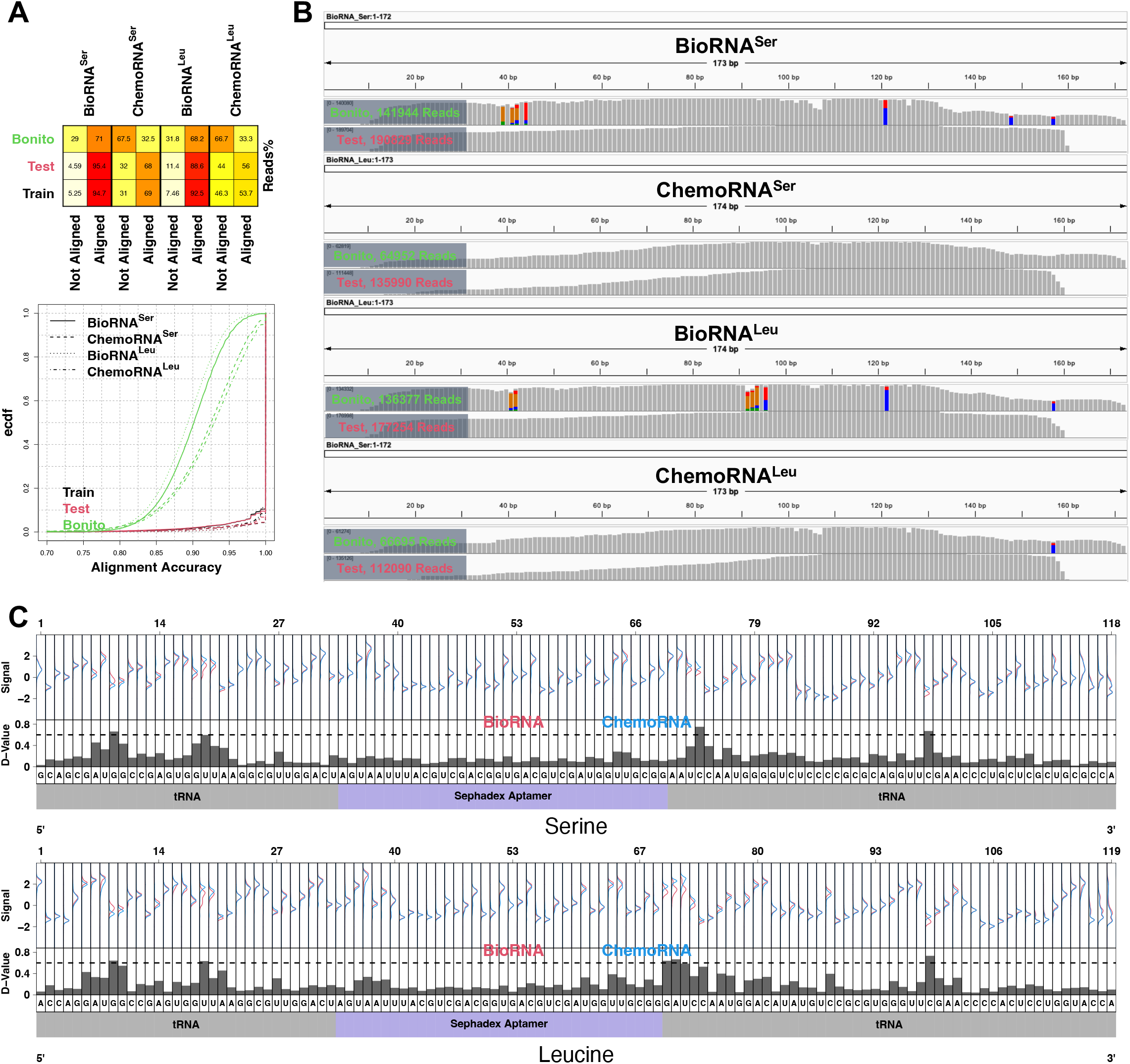
Determining BioRNA modification status based on iteratively basecalled sequence backbones. (A) Mappability and alignment accuracy of BioRNAs (containing modifications) and ChemoRNAs (no modification). Train, polished basecalling results for datasets used during iterative basecaller training; Test, polished basecalling results for independent test datasets; Bonito, initial Bonito basecalling results for test datasets. (B) IGV visualization for initial Bonito and polished basecalling results. (C) The sequencing signal comparison between BioRNAs and corresponding ChemoRNAs. Signals mapped to the same nucleotide position were summarized using a probability density distribution (upper panel). Distribution differences between Bio and ChemoRNAs were quantified by KS-test D-values (lower panel): larger D-values indicate higher modification likelihood.

## DISCUSSION

In nanopore sequencing, precisely resolving sequence backbones is the prerequisite for virtually all the downstream bioinformatic analyses. In this paper, we present an iterative approach, which could polish basecalling results, based on our *a priori* knowledge about the ground-truth reference sequence. Our approach is thus uniquely suitable for training biomolecule-specific high-accuracy basecallers. For instance, we confirmed the superior performance in basecalling control oligos, as well as native rRNAs and tRNAs using our approach. Building upon such benchmarked basecalling performance, we leveraged our approach for the quality control of therapeutic molecules. Specifically, we basecalled the VAX-seq data as the basis for quantifying the mRNA vaccine purity and integrity, as well as the BioRNA data as the basis for inspecting modification hotspots in RNAi drugs.

The success of our iterative approach depends on the performance of initial basecalling. As a result, “non-basecallable” nanopore sequencing signals, such as those significantly deviated by modifications, are unlikely to be properly handled, as one major limitation of our approach. We therefore expect the development of generic basecallers, in particular ones that are tolerant to various categories of modifications as described in our previous research^18^, to be one potential solution.

During the iterative basecalling process, sequence backbones will be polished based on the reference sequence. Thus, sequence discrepancies between the provided reference and real-world biomolecules could introduce systematic basecalling artifacts, as another limitation of our approach. For instance, single nucleotide polymorphism sites, which are generally not recorded in the reference, cannot be properly handled by our approach. To maximize the performance of iterative basecalling, the quality of reference sequence, as well as the sequence homogeneity of candidate biomolecules, need to be guaranteed.

While this paper primarily focuses on the polishing of modification-disturbed basecalling, our iterative approach could also promote the interpretation of canonical sequences. For example, basecalling accuracy of unmodified RNAs, such as rRNAs produced from IVT, tRNA species Gln-CTG and chemically-synthesized ChemoRNAs, was largely improved by iterative basecalling (Figure 2 and 4). Therefore, we highlight the iterative basecalling as a general approach for polishing sequence backbones.

## METHODS

### The Production and Purification of BioRNAs

The design and production of bioengineered human seryl-tRNA (TGA) and leucyl-tRNA (TAA) (BioRNA^Ser^ and BioRNA^Leu^) have been described in our previous studies^19,20^. In brief, human tRNA sequences were obtained from GtRNAdb (https://gtrnadb.ucsc.edu/) and anti-codon regions were replaced by a Sephadex aptamer to establish recombinant human tRNA constructs. Target tRNA coding sequences (Table S1) were amplified with polymerase chain reaction (PCR) with specific primers. RNA expression plasmids were then constructed by infusing amplicons into a pBSTNAV vector with the In-Fusion® HD Cloning kit (Takara). After verification with DNA sequencing, plasmids were transformed to *E. coli* HST08 competent cells to express and accumulate BioRNA^Ser^ and BioRNA^Leu^ through overnight fermentation on a large scale (∼500 mL). The total RNA was isolated from the overnight culture using the phenol extraction method, and subsequently loaded and separated on the ENrich-Q 10×100 anion exchange column (Bio-Rad) integrated in the NGC Quest 10 Plus chromatography system (Bio-Rad) for purifying target BioRNAs. The total RNA was eluted using the gradient method, which consist of buffer A (10 mM monosodium phosphate, pH 7.0) and buffer B (buffer A + 1 M sodium chloride, pH 7.0) at a constant flow rate of 2 mL/min. To purify BioRNAs, the column was first equilibrated with buffer A for 6.7 min, followed by elution with 55% buffer B for 4.8 min, then 55-70% buffer B for 40 min (for BioRNA^Ser^), or 55-75% buffer B for 30 min (for BioRNA^Leu^). The column was subsequently washed using 100% buffer B for 10 min, with an additional re-equilibration using buffer A for 10 min. Fractions of target RNA peaks were collected during the total RNA elution, and then loaded onto urea PAGE gels to verify size and homogeneity. Pure fractions were combined, desalted and concentrated using Amicon Ultra-2mL centrifugal filters (30 k; MilliporeSigma) to obtain ready-to-use pure BioRNAs. The BioRNA purity was determined by the urea PAGE analysis, and quantified using the high-performance liquid chromatography (HPLC) as previously reported^20,21^. BioRNA products with high homogeneity (> 98%) were used in the nanopore sequencing study.

### ChemoRNAs and Nano-tRNAseq Adaptors

The chemically synthesized, Sephadex aptamer-tagged human seryl-tRNA (TGA) and leucyl-tRNA (TAA) (ChemoRNA^Ser^ and ChemoRNA^Leu^), and Nano-tRNAseq^9^ adapters were ordered from Integrated DNA Technologies. Specifically, for nano-tRNAseq 5’ RNA splint adapters, the first four nucleotides at the 3’ terminal were designed as rUrGrGrC and rUrGrGrU, which are complementary to seryl-tRNA and leucyl-tRNA, respectively.

Other Nano-tRNAseq adapters, including 3’ RNA:DNA splint adapters and ONT reverse transcription adapters (RTAs, oligo A and oligo B) were synthesized as reported in the original study. Sequences of ChemoRNAs and adapters were listed in Table S1.

### The Nanopore Sequencing of BioRNAs and ChemoRNAs

BioRNA and ChemoRNA nanopore sequencing libraries were constructed first following the nano-tRNAseq protocol^9^, then using the Direct RNA Sequencing Kit (SQK-RNA004, Oxford Nanopore Technologies) following manufacturer’s instructions. Specifically:

#### Adapter Annealing

5’ RNA splint adapters and equal moles (∼108 pmol) of 3’ RNA:DNA splint adapters were mixed with a dilution buffer consisting of Tris-HCl (10 mM, pH 7.5), NaCl (500 mM), and 1 μL of RNasin Ribonuclease Inhibitor (N2511, Promega) to a final concentration of ∼50 ng/μL for each adapter in a total volume of 20 μL. The mixture was incubated at 75 °C for 15 seconds, further cooled to 25 °C at a rate of 0.1 °C/s (a total of 500 seconds) to anneal splint adapters. RTA oligo As and oligo Bs (∼82 pmol for each, ∼200 ng in total) were annealed under the same condition as splint adapters.

#### Ligation 1

1 μg of target RNA (ChemoRNA^Ser^, ChemoRNA^Leu^, BioRNA^Ser^ or BioRNA^Leu^) was ligated to pre-annealed splint adapters (RNA:adapters = 1.2:1, molar ratio) in a 20 μL reaction system containing T4 RNA ligase 2 (10U, M0239S, Promega), 1 × Reaction Buffer (B0216S, NEB), 10% PEG 8000 (B0216S), 400 μM ATP (B0216S), and 1 µL RNasin Ribonuclease Inhibitor. The reaction was conducted at room temperature for 1 hour. Ligation products were purified with 1.8x volume of well-mixed, room temperature AMPure RNAClean XP beads (A63987, Beckman Coulter) following manufacturer’s instructions. Concentrations of purified samples were determined using the Tecan plate reader (Tecan), and the quality of ligated RNAs was evaluated by urea polyacrylamide gel electrophoresis (PAGE) analyses.

#### Ligation 2 and reverse transcription (RT)

Purified “Ligation 1” products (400 ng, ∼10.4 pmol, for two sequencing replicates) were further ligated to pre-annealed RTA adapters (RNA:adapters = 1:2, molar ratio) with the T4 DNA Ligase (M0202M, NEB) in a 30 μL reaction buffer constituted by 6 μL of NEBNext Quick Ligation Reaction Buffer (B6058S, NEB), 1 μL of RNasin Ribonuclease Inhibitor and RNase-free water by incubating at room temperature for 30 min. Subsequently, the ligation product was mixed with 4 μL of dNTPs (10 mM, N0447S, NEB) and 26 μL of RNase-free water to pre-denature the RNA at 65 °C for 5 min and then cooled down on ice for 2 min. A mixture of Maxima H Minus Reverse Transcriptase (4 μL, EP0751, Thermo Fisher Scientific), 16 μL of Maxima H Minus Reverse Transcriptase Buffer, and 2μL of RNasin Ribonuclease Inhibitor (80U) were added to the pre-treated RNA sample to perform RT by incubation at 60 °C for 1 hour, 85 °C for 5 min, then cooling to 4 °C. AMPure RNAClean XP beads were then added to purify the RT products according to the protocol. The quantity and quality of the purified samples were determined by the Tecan plate reader and the urea PAGE gel, respectively.

#### Ligation 3 and final library

Following purification, 40 μL (two replicates) of RT products were ligated to 12 μL of RNA Ligation Adapter (RLA, provided by the SQK-RNA004 kit) in an 80 μL reaction system, supplemented with 16 μL of NEBNext Quick Ligation Reaction Buffer and 6 μL of T4 DNA Ligase for a 30-min incubation at room temperature. The ligation products were purified using AMPure RNAClean XP beads following the protocol provided by the SQK-RNA004 kit. Afterwards, MinION flow cells (RNA chemistry) were primed following manufacturer’s instructions. Meanwhile, Elution Buffer (provided by the SQK-RNA004 kit) re-suspended libraries were gently mixed with 37.5 μL of Sequencing Buffer (provided by the SQK-RNA004 kit) and 25.5 μL of Library Solution (provided by the SQK-RNA004 kit) to obtain final libraries. A total volume of 75 μL final library was loaded to each flow cell for sequencing.

The step-by-step validation of RNA purity and quality throughout the library preparation process was presented in Figure S4.

### Iterative Basecalling with the Guppy-Taiyaki Workflow

RNA control oligos, yeast rRNAs and tRNAs, as well as vaccine mRNAs were iteratively basecalled with the Guppy-Taiyaki workflow. The workflow consists of 1) Guppy (version 6.0.6+8a98bbc) for basecalling and alignment; 2) Samtools (version 1.16) for alignment result processing including merge, sort and index^22^; 3) Taiyaki (version 5.3.0) for training Guppy model, including 3.1) data preparation by scripts generate_per_read_params.py, get_refs_from_sam.py, prepare_mapped_reads.py and merge_mappedsignalfiles.py; 3.2) model training by scripts train_flipflop.py and dump_json.py; 4) iterating the above steps with the trained model. The workflow is summarized in Figure S5.

For Step 1, the initial basecalling used Guppy model “template_r9.4.1_450bps_hac.jsn”, together with the “--disable_qscore_filtering” flag to keep as many reads as possible for downstream analyses. All other flags were set default. Subsequent iterations used the trained model with the “--disable_qscore_filtering” flag, and other flags were set default. In particular, a stand-alone Minimap2^23^ (version 2.24-r1122) was used to optimize tRNA alignments, with “-ax map-ont -k5 -w5” flags as recommended in the original study^9^.

For Step 2, default flags for Samtools merge, sort and index were used for all analyses.

For Step 3, The “--reverse” flag was used for get_refs_from_sam.py, with other flags set default. Flags for generate_per_read_params.py and merge_mappedsignalfiles.py were set default. The initial iteration used the model checkpoint “r941_rna_minion.checkpoint” that is provided by Taiyaki, to run the prepare_mapped_reads.py (a customized version: https://github.com/wangziyuan66/IL-AD/blob/main/scripts/trna/train_flipflop.py was used for the tRNA analysis); the model checkpoint trained in the previous iteration was used for subsequent iterations. All other flags were set default during this process.

As for model training, the model template “mLstm_flipflop.py” that is provided by Taiyaki, and flags “--size 256 --stride 10 --winlen 31”, were used for train_flipflop.py. Other flags were set default. Default flags for dump_json.py were used for generating final models.

### Iterative Basecalling with the Bonito Workflow

BioRNAs and ChemoRNAs were iteratively basecalled with the Bonito (version 0.8.1) workflow, including 1) the “bonito baseball” option for basecalling, alignment and training data preparation; 2) the “bonito train” option for model training; 3) iterating Step 1 and 2 with the trained model. The workflow is summarized in Figure S6.

For Step 1, the initial basecalling used Bonito model “rna004_130bps_hac@v3.0” with “--chunksize 3000 --save-ctc --min-accuracy-save-ctc 0.9” flags to prepare high-quality training data. All other flags were set default. Subsequent iterations used the trained model with flags “--chunksize 3000 –save-ctc --min-accuracy-save-ctc 0.9”, and other flags were set default.

For Step 2, the initial model training fine-tunes the model “rna004_130bps_hac@v3.0”. Subsequent iterations fine-tune the model produced from the previous iteration. Flags “--size 256 --epochs 5 --lr 5r-4” (other flags were set default) during the training process.

### Basecalling Performance Evaluation

Basecalling performance was “functionally” evaluated by downstream alignment results, including mappability (the ratio between aligned reads, regardless of SAM flags, and all reads), alignment accuracy (ratio between CIGAR M, and CIGAR M and D; please note that Minimap2 represents X as M in CIGAR when aligning nanopore sequencing reads), alignment consistency (alignment start and end positions) as opposed to Guppy results, and IGV visualization (version 2.17.4). Specifically, for RNA control oligos, yeast rRNAs and vaccine mRNAs, alignments results were directly obtained from Guppy outputs. For yeast tRNAs and Bio/ChemoRNAs, a stand-alone Minimap2^23^ (version 2.24-r1122) was used for alignment with “-ax map-ont -k5 -w5” flags as previously recommended^9^.

### Nanopore Sequencing Signal Analysis

We first basecalled nanopore sequencing reads using Bonito (version 0.8.1) to produce move-tables, which track signal chunks corresponding to individual nucleotides (stored as mv tags in the generated bam files). We next executed the Remora (version 3.2.0) pipeline “Reference Region Metric Extraction” (https://github.com/nanoporetech/remora/blob/master/notebooks/metrics_api.ipynb) to extract the per-nucleotide signal mean values. Throughout the analysis, all the Bonito parameters were set as default. The “reverse_signal” flag for Remora was set as True, and all the other Remora parameters were set as default.

### Data Availability

The BioRNA nanopore sequencing data was deposited at NCBI under the BioProject PRJNA1155679. The RNA oligo nanopore sequencing data was downloaded from NCBI under the BioProject PRJNA1050579 (runs SRR27324841, SRR27324838 and SRR27324839). Corresponding reference sequences were downloaded from the Table S2 of ^8^. The yeast native rRNA nanopore sequencing data was downloaded from ENA under accession number PRJEB48183 (samples ERR7162388, ERR7162390 and ERR7162391). Corresponding reference sequences were downloaded from https://github.com/adbailey4/yeast_rrna_modification_detection/tree/main/notebooks/data. The yeast native tRNA nanopore sequencing data was downloaded from ENA under accession number PRJEB55684 (samples ERR10224971 and ERR10224975). Corresponding reference sequences and modification annotations were downloaded from https://github.com/novoalab/Nano-tRNAseq/tree/main/ref. The mRNA vaccine nanopore sequencing data was downloaded from NCBI under the BioProject PRJNA856796 (runs SRR22888949 and SRR22888950). Corresponding reference sequence was downloaded from https://github.com/scchess/Mana/tree/main/data.

### Code Availability

The Taiyaki iterative basecalling framework can be found at: https://github.com/wangziyuan66/iterative-labeling-toolkit-taiyaki. The Bonito iterative basecalling framework can be found at: https://github.com/wangziyuan66/iterative-labeling-toolkit-bonito.

## ACKNOWLEDGEMENTS

We thank Dr. Xinlei Sheng for his valuable discussions. We thank the University of Arizona High Performance Computing team and the College of Pharmacy Information Technology Group for their support. H.D. is supported by the University of Arizona Health Sciences Career Development Award, and the University of Arizona Accelerate For Success Award. A.-M.Y. is supported by the National Institute of General Medical Sciences [R35GM140835] and National Cancer Institute [R01CA225958 and R01CA253230], National Institutes of Health (NIH). J.Q. is supported by the National Heart, Lung, and Blood Institute [R01HL159675 and R01HL152293], National Institute of Allergy and Infectious Diseases [R21AI163753] and National Institute of Diabetes and Digestive and Kidney Diseases [R01DK132251], National Institutes of Health (NIH).

## AUTHOR CONTRIBUTIONS

H.D. and A.-M.Y. conceived the idea. Z.W., Z.L. and H.D. performed the analysis. M.-J.T., K.K.W. and Y.F. performed the experiment. N.H., H.H.Z., X.S., J.Q., A.-M.Y. and H.D. supervised the project. Z.W., M.-J.T., A.-M.Y. and H.D. wrote the manuscript.

