## Supplementary for "An Iterative Approach to Polish the Nanopore Sequencing Basecalling for Therapeutic RNA Quality Control"

### Supplementary Information

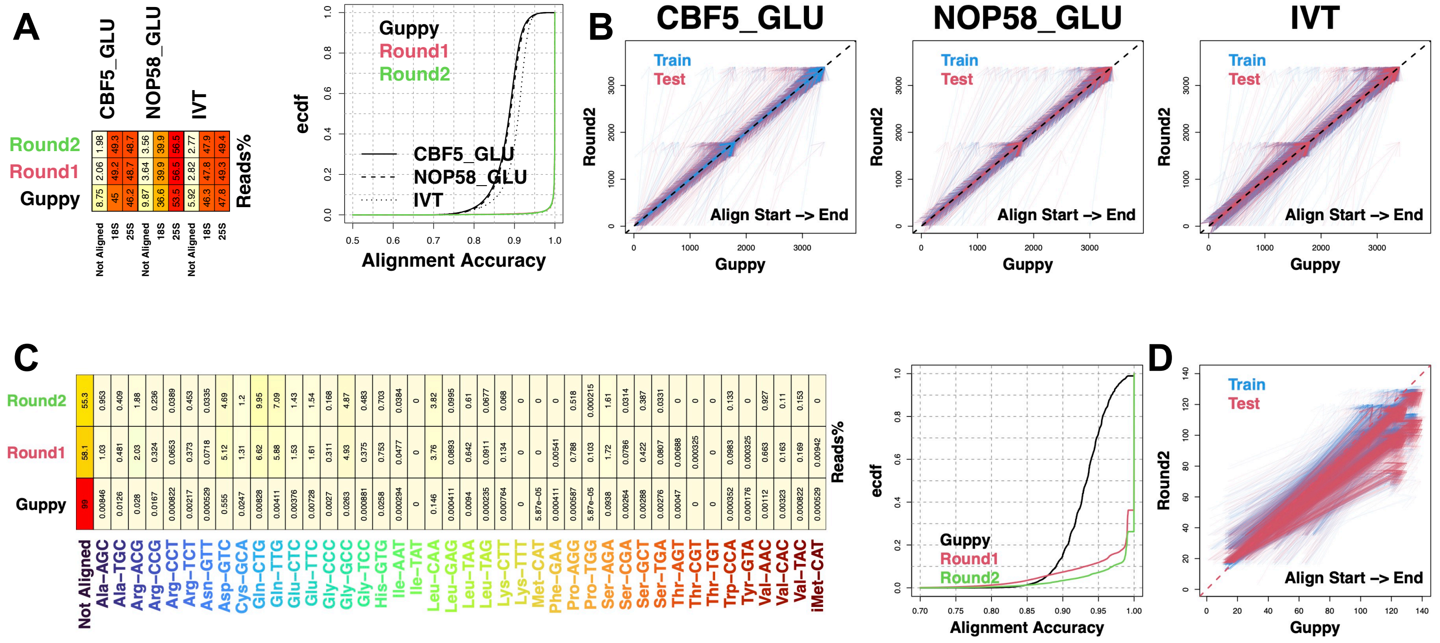

A
CBF5_GLU
NOP58_GLU
IVT
Round2
Round1
Guppy
Reads%
Not Aligned
18S
25S
1.98 49.3 48.7 3.56 39.9 56.5 2.77 47.9 49.4
2.06 49.2 48.7 3.64 39.9 56.5 2.82 47.8 49.3
8.75 45 46.2 9.87 36.6 53.5 5.92 46.3 47.8
ecdf
Alignment Accuracy
Guppy
Round1
Round2
CBF5_GLU
NOP58_GLU
IVT
B
CBF5_GLU
NOP58_GLU
IVT
Train
Test
Round2
Guppy
Align Start -> End
C
Round2
Round1
Guppy
Reads%
Not Aligned
Ala-AGC
Ala-TGC
Arg-ACG
Arg-CCG
Arg-CCT
Arg-TCT
Asn-GTT
Asp-GTC
Cys-GCA
Gln-CTG
Gln-TTG
Glu-CTC
Glu-TTC
Gly-CCC
Gly-GCC
Gly-TCC
His-GTG
Ile-AAT
Ile-TAT
Leu-CAA
Leu-GAG
Leu-TAA
Leu-TAG
Lys-CTT
Lys-TTT
Met-CAT
Phe-GAA
Pro-AGG
Pro-TGG
Ser-AGA
Ser-CGA
Ser-GCT
Ser-TGA
Thr-AGT
Thr-CGT
Thr-TGT
Trp-CCA
Tyr-GTA
Val-AAC
Val-CAC
Val-TAC
iMet-CAT
ecdf
Alignment Accuracy
Guppy
Round1
Round2
D
Train
Test
Round2
Guppy
Align Start -> End

**Figure S1. Training and benchmarking iterative basecallers for the real-world RNA molecule sequence analysis.** (A) Yeast rRNA mappability and alignment accuracy. Distributions of alignment accuracy were shown with ecdf (empirical cumulative distribution function). CBF5_GLU and NOP58_GLU, mutant strain rRNAs which have different RNA modification patterns; IVT, rRNAs made by *in vitro* transcription which contain no modification. Guppy, Guppy basecalling results; Round1 & 2; basecalling results produced by subsequent iterations. (B) Alignment position comparison between Guppy and Round2 results. Train, results for train datasets used in panel A; Test, results for independent test datasets. Arrows represent alignment directions. (C) Yeast native tRNA mappability and alignment accuracy. Distributions of alignment accuracy were shown with ecdf (empirical cumulative distribution function). Guppy, Guppy basecalling results; Round1 & 2; basecalling results produced by subsequent iterations. (D) Alignment position comparison between Guppy and Round2 results. Train, results for train datasets used in panel C; Test, results for independent test datasets. Arrows represent alignment directions.

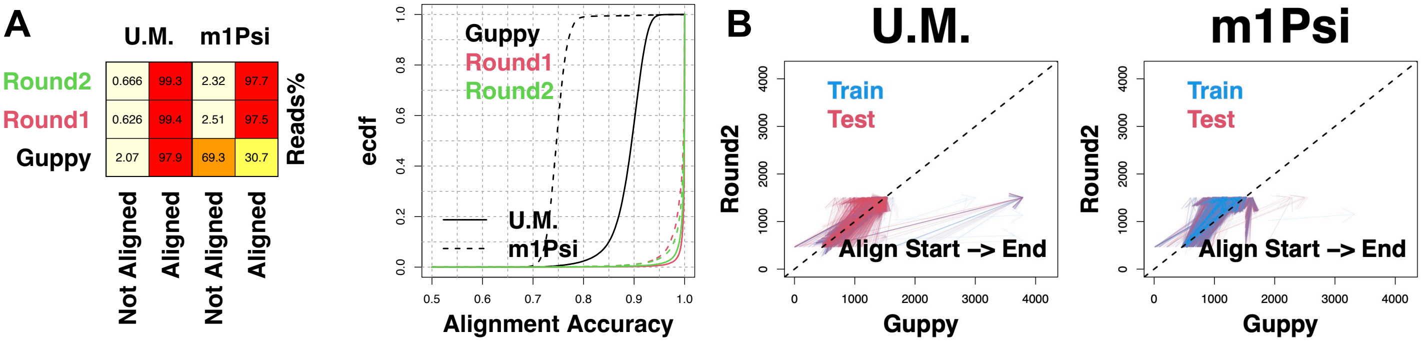
A
U.M.
m1Psi
Round2
Round1
Guppy
0.666
99.3
2.32
97.7
0.626
99.4
2.51
97.5
2.07
97.9
69.3
30.7
Reads%
Not Aligned
Aligned
Not Aligned
Aligned
Guppy
Round1
Round2
U.M.
m1Psi
ecdf
Alignment Accuracy
B
U.M.
m1Psi
Train
Test
Round2
Guppy
Align Start –> End

**Figure S2. Training and benchmarking iterative basecallers for the vaccine mRNA sequence analysis.** (A) Mappability and alignment accuracy. Distributions of alignment accuracy were shown with ecdf (empirical cumulative distribution function). U.M., canonical mRNA; m1Psi, U-to-N1-methylpseudouridines fully-replaced mRNA. Guppy, Guppy basecalling results; Round1 & 2; basecalling results produced by subsequent iterations. (B) Alignment position comparison between Guppy and Round2 results. Train, results for train datasets used in panel A; Test, results for independent test datasets. Arrows represent alignment directions.

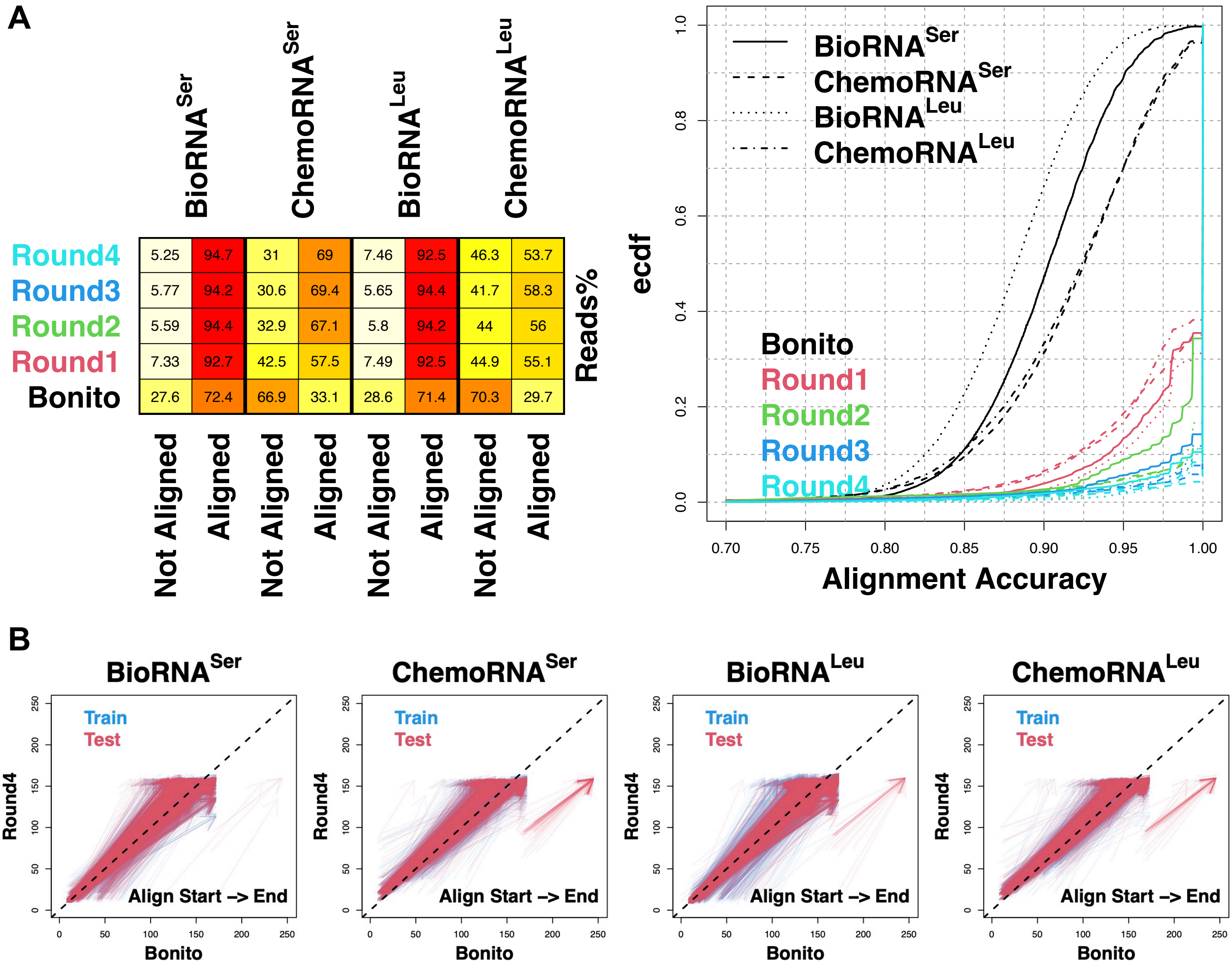
A
BioRNA^Ser
ChemoRNA^Ser
BioRNA^Leu
ChemoRNA^Leu
Round4 5.25 94.7 31 69 7.46 92.5 46.3 53.7
Round3 5.77 94.2 30.6 69.4 5.65 94.4 41.7 58.3
Round2 5.59 94.4 32.9 67.1 5.8 94.2 44 56
Round1 7.33 92.7 42.5 57.5 7.49 92.5 44.9 55.1
Bonito 27.6 72.4 66.9 33.1 28.6 71.4 70.3 29.7
Not Aligned
Aligned
Reads%
ecdf
Alignment Accuracy
BioRNA^Ser
ChemoRNA^Ser
BioRNA^Leu
ChemoRNA^Leu
Bonito
Round1
Round2
Round3
Round4
B
BioRNA^Ser
ChemoRNA^Ser
BioRNA^Leu
ChemoRNA^Leu
Train
Test
Align Start -> End
Round4
Bonito

**Figure S3. Training and benchmarking iterative basecallers for the BioRNA and ChemoRNA sequence analysis.** (A) Mappability and alignment accuracy. Distributions of alignment accuracy were shown with ecdf (empirical cumulative distribution function). Bonito, Bonito basecalling results; Round1-4; basecalling results produced by subsequent iterations. (B) Alignment position comparison between Bonito and Round4 results. Train, results for train datasets used in panel A; Test, results for independent test datasets. Arrows represent alignment directions.

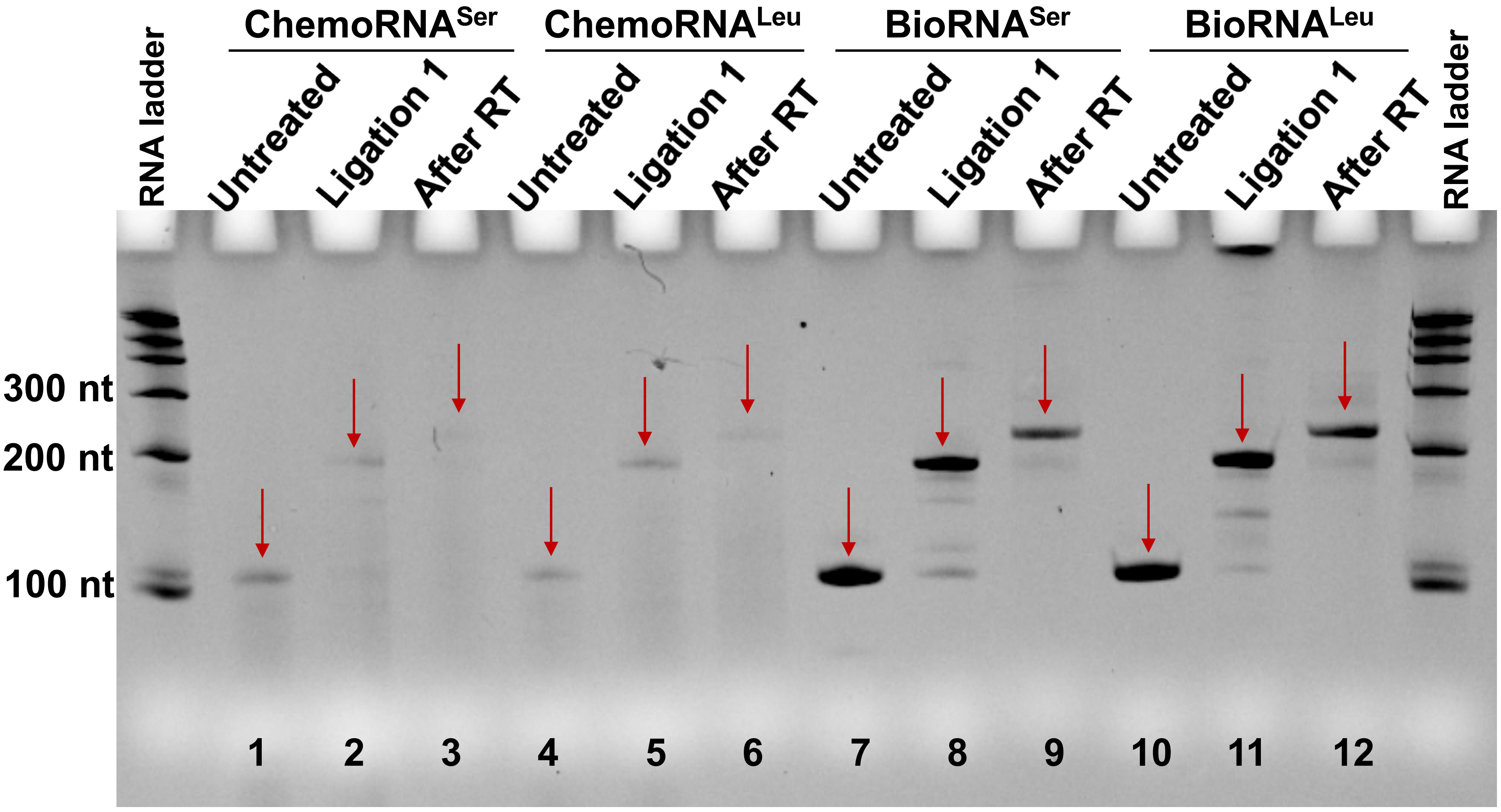
ChemoRNA$^{Ser}$
ChemoRNA$^{Leu}$
BioRNA$^{Ser}$
BioRNA$^{Leu}$
RNA ladder
Untreated
Ligation 1
After RT
Untreated
Ligation 1
After RT
Untreated
Ligation 1
After RT
Untreated
Ligation 1
After RT
RNA ladder
300 nt
200 nt
100 nt
1
2
3
4
5
6
7
8
9
10
11
12

**Figure S4. The step-by-step validation of RNA purity and quality throughout the Nano-tRNAseq nanopore sequencing library preparation process.** Denaturing urea polyacrylamide gel electrophoresis (PAGE) was used to analyze the purity of candidate RNAs (ChemoRNA$^{Ser}$, ChemoRNA$^{Leu}$, BioRNA$^{Ser}$, BioRNA$^{Leu}$) and their ligation products. Lanes 1 and 4: ChemoRNA$^{Ser}$ (118 nt) and ChemoRNA$^{Leu}$ (119 nt) showed visible bands at expected sizes, together with impurity signals. These results indicated compromised purity of chemically synthesized commercial products. Lanes 7 and 10: BioRNA$^{Ser}$ (118 nt) and BioRNA$^{Leu}$ (119 nt) exhibited unique and distinct bands at expected sizes. These results indicated the high homogeneity of BioRNAs produced using our bioengineering system. Lanes 2, 5, 8 and 11: Purified products after the 5' and 3' splint adapter ligation (Ligation 1, 57 nt were added). Lanes 3, 6, 9 and 12: Purified products after the reverse transcription adapter (RTA) ligation (Ligation 2) and the reverse transcription procedure (RT). Abundant band signals were observed at the expected positions for all the ligation and RT products, with moderate intensity of by-products that could not be removed with purification. For the PAGE analysis, the sample loading quantity was 100 ng/lane.

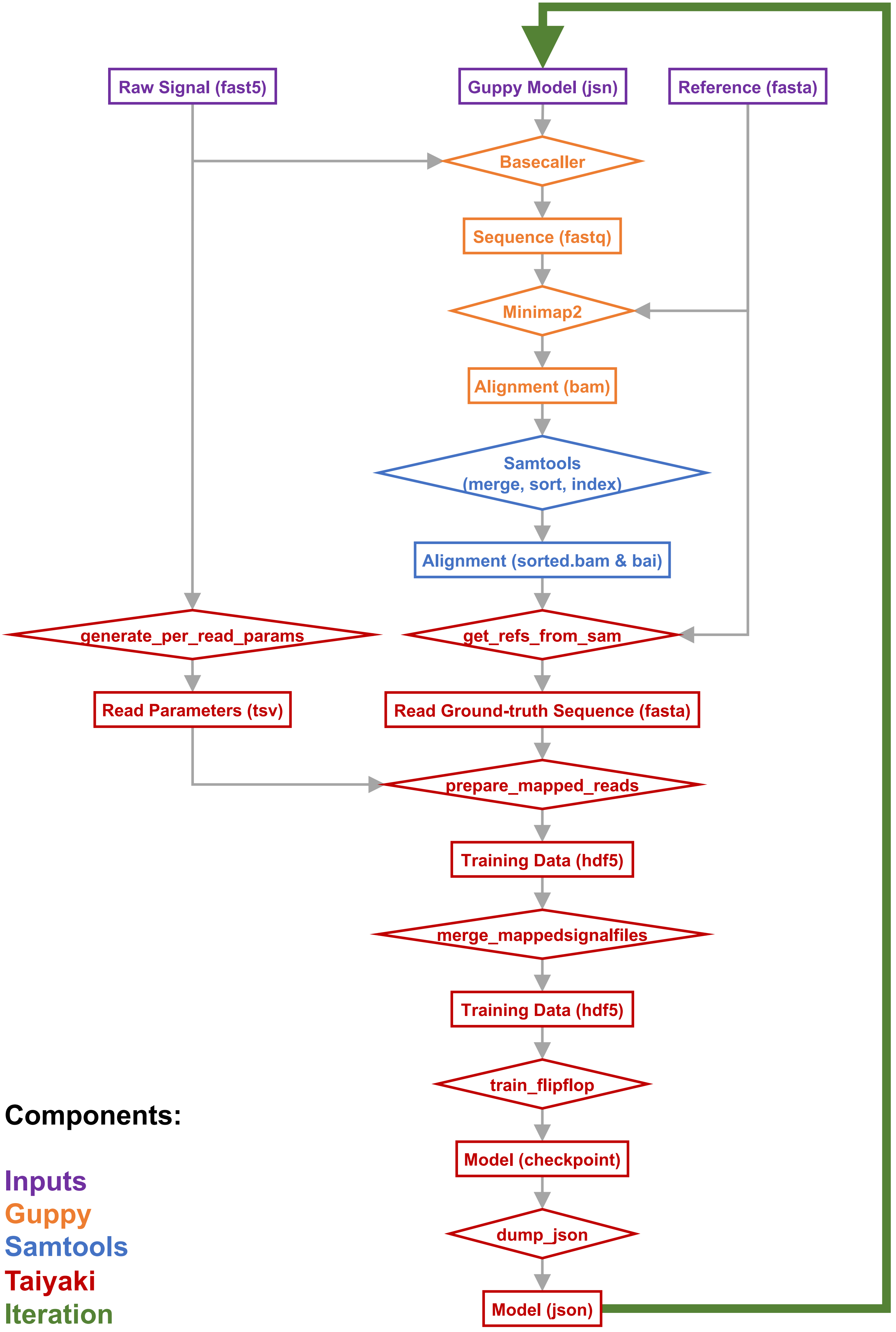
Raw Signal (fast5)
Guppy Model (jsn)
Reference (fasta)
Basecaller
Sequence (fastq)
Minimap2
Alignment (bam)
Samtools
(merge, sort, index)
Alignment (sorted.bam & bai)
generate_per_read_params
get_refs_from_sam
Read Parameters (tsv)
Read Ground-truth Sequence (fasta)
prepare_mapped_reads
Training Data (hdf5)
merge_mappedsignalfiles
Training Data (hdf5)
train_flipflop
Model (checkpoint)
dump_json
Model (json)
Components:
Inputs
Guppy
Samtools
Taiyaki
Iteration

**Figure S5. Guppy-Taiyaki iterative basecalling workflow.**

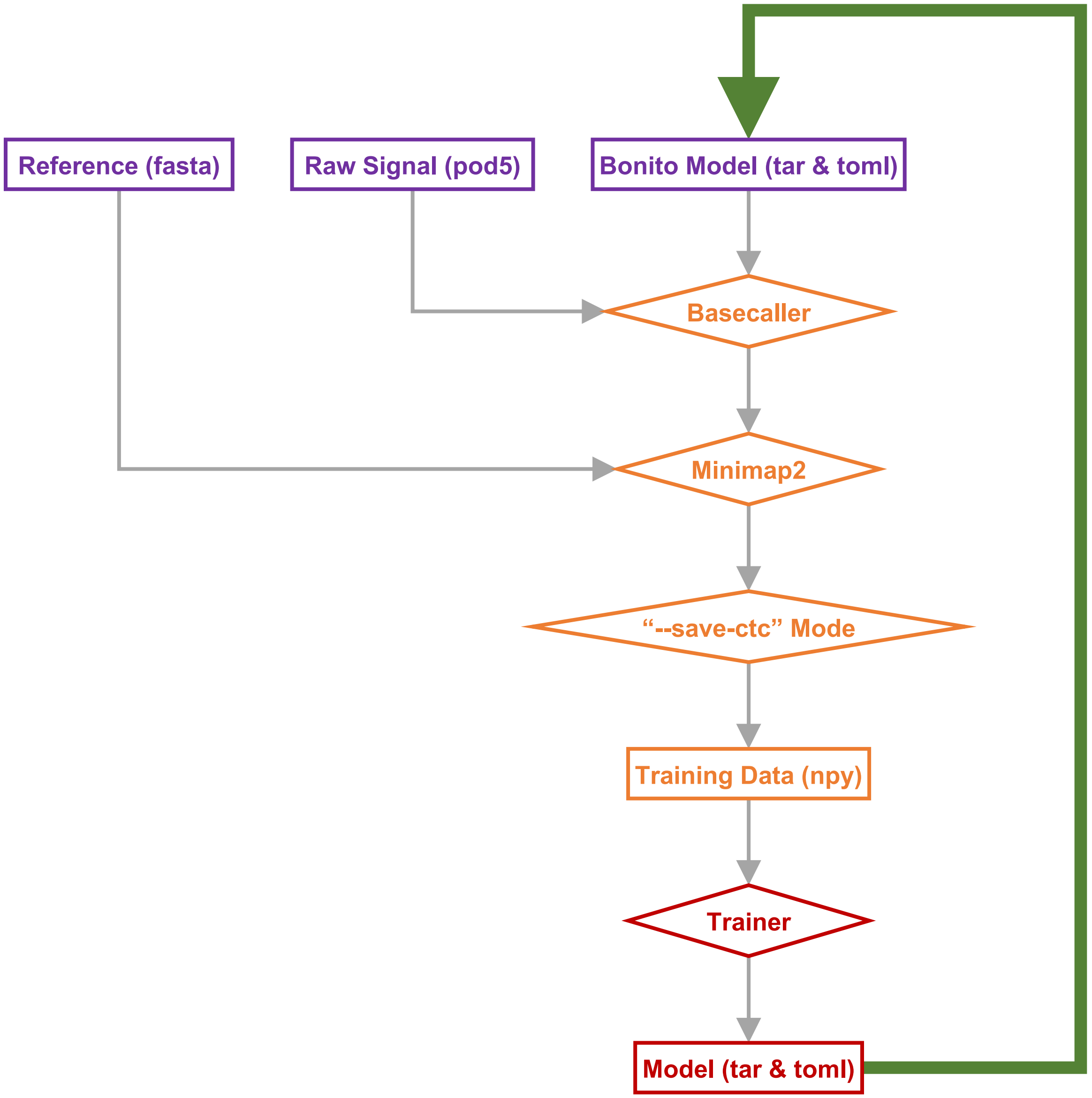


**Components:**

Inputs
Bonito Basecaller
Bonito Trainer
Iteration

**Figure S6. Bonito iterative basecalling workflow.**

**Table S1. Sequences of Bio/ChemoRNAs and adapters.**
